# Development and Application of a Tri-allelic PCR Assay for Screening *Vgsc*-L1014F *Kdr* Mutations Associated with Pyrethroid and Organochlorine Resistance in the Mosquito *Culex quinquefasciatus*

**DOI:** 10.1101/588905

**Authors:** Walter Fabricio Silva Martins, Bárbara Natieli Silva Pereira, Ana Thayse Vieira Alves, Annabel Murphy, Paulo Geovani Silva Martins, David Weetman, Craig Stephen Wilding, Martin James Donnelly

## Abstract

**Background:** *Culex quinquefasciatus,* has a widespread distribution across tropical and sub-tropical regions, and plays an important role in the transmission of vector-borne diseases of public health importance, including lymphatic filariasis (LF) and multiple arboviruses. Increased resistance to insecticides threatens the efficacy and sustainability of insecticide-based anti-vector interventions which mitigate the burden of mosquito transmitted diseases in endemic regions. In *C. quinquefasciatus* two non-synonymous voltage gated sodium channel (*Vgsc*) variants, both resulting in a leucine to phenylalanine change at codon 1014, are associated with resistance to pyrethroids and DDT. This tri-allelic variation has compromised the ability to perform high-throughput single-assay screening. To facilitate the detection and monitoring of the *Vgsc*-1014 locus in field-caught mosquitoes, an Engineered-Tail Allele-Specific-PCR (ETAS-PCR) diagnostic assay was developed and applied to wild mosquitoes from Brazil, Tanzania and Uganda.

**Results:** This new cost-effective, single-tube assay was compared to two, well-established, genotyping approaches – pyrosequencing and TaqMan. The ETAS-PCR assay showed high specificity for discriminating the three alleles at *Vgsc*-L1014F, with genotyping results strongly correlated with 98.64% and 100% against pyrosequencing and TaqMan, respectively.

**Conclusions:** Our results support the utility of the ETAS-PCR/*Vgsc*-1014 diagnostic assay, which stands as an effective alternative for genotyping tri-allelic variants.

## Background

The mosquito *Culex pipiens quinquefasciatus* (hereafter *C. quinquefasciatus*) acts as a vector of several pathogens in both tropical and temperate environments (1). In many tropical/sub-tropical regions *C. quinquefasciatus* is the primary vector of lymphatic filariasis (LF) (2-4) whilst in subtropical locations, it is a vector of potentially fatal arboviruses e.g. West Nile Virus (WNV) and St. Louis encephalitis Virus (SLEV) (5-7).

Vaccination and mass drug administration (MDA) are the primary methods for reducing the burden of diseases transmitted by *C. quinquefasciatus,* but insecticide-based vector control is advocated as an important adjunct. For example, vector control has assisted in mitigating the burden of transmission of arboviruses where immunization campaigns are challenging, such as in rural settlements (8-10).

The effectiveness of insecticide-based vector control is threatened by the increased insecticide resistance detected across the globe (11, 12) with the increased resistance to pyrethroids especially worrying as it is the most common active ingredient used to control adult mosquitoes through indoor sprays, outdoor fogs and treated bednets (13).

In *C. quinquefasciatus* as well as in other mosquitoes of public health importance pyrethroid and DDT-resistance has been associated with two major mechanisms; overexpression of detoxification genes (14-16) and a variety of alleles in the Voltage-Gated Sodium Channel (*Vgsc*) gene – called knockdown resistance (*kdr*) mutations such as the *Vgsc*-1014F (17-19). Consequently, developing molecular tools to detect and monitor resistance-alleles at this locus are imperative for the study of the evolution of resistance, and may also assist programme managers in the rational deployment of insecticides.

Several diagnostic tests for typing *kdr* mutations have already been developed and applied in *Culex, Anopheles* and *Aedes* species (20-22). Whilst many rely on standard PCR methods (23-26), over the past few years there has been an increase in the use of high-throughput methods such as quantitative PCR (e.g. TaqMan allele-specific and melt curve analysis) and pyrosequencing (18, 21, 27).

Regardless of methodology, most assays are designed to perform bi-allelic discrimination, and there are a limited number of low cost approaches for typing tri-allelic mutations (e.g. (20, 24, 28) such as the alternative codons (TTA/TTT/TTC) at locus *Vgsc*-1014 already reported in *C. quinquefasciatus* populations from Africa and Asia (18, 21, 29).

To address this limitation, we designed and applied a new diagnostic assay – ETAS-PCR (Engineered-Tail Allele-Specific-PCR), which runs under standard PCR conditions, to type tri-allelic variation at *Vgsc-*1014. Samples of field-caught mosquitoes from Brazil, Tanzania and Uganda were genotyped using the ETAS-PCR/*Vgsc*-1014 assay along with pyrosequencing and TaqMan allele-discrimination to assess concordance.

## Materials and methods

### Ethics Statement

No specific permits were required for the described field work as no human participants were involved. Oral consent was obtained from householders prior to collections and no personal details were recorded.

### Sampling

A total of 183 field-caught and pyrethroid-or DDT-selected *C. quinquefasciatus* individuals from Brazil, Tanzania and Uganda were genotyped. Mosquitoes from Campina Grande (07° 13′ 50″ S, 35° 52′ 52″ W), Paraíba, Brazil were collected in 2016 using 50 ovitraps, one per dwelling randomly distributed within a single neighbourhood. Blood-fed mosquitoes from Tanzanian, Mwanza (02° 28’ S, 32° 55’ E) were collected from inside houses using a manual aspirator in 2013. Field-caught mosquitoes from Tororo (0°40′41.62″N, 34°11′11.64″E) and Jinja (0° 26′ 52.285″ N 33° 12′ 9.403″ E), Uganda, were collected in 2012 while Tororo insecticide exposed mosquitoes were phenotyped using insecticide bioassays as described previously (14, 21). Additionally, 12 specimens from Uganda, genotyped by Sanger sequencing were used as a control (27).

Genomic DNA of individual mosquitoes was isolated using the DNeasy Kit (Qiagen) and samples were confirmed as *C. quinquefasciatus* through a diagnostic PCR assay (30).

### ETAS-PCR Assay design

#### Primer Design

Genetic variation in primer-binding sites could result in null alleles and thereby erroneous PCR genotyping. Therefore, conserved regions in the vicinity of the *Vgsc*-1014 locus were selected from an alignment of 10 *C. quinquefasciatus* sequences from distinct geographic regions (Additional file 1. Figure S1), and 6 sequences from other *Culex pipiens* sub-species: *C. pipiens quinquefasciatus*, Saudi Arabia, Thailand, Uganda and USA and *C. pipiens pallens* and *C. pipiens pipiens* from China and USA, respectively (see Additional file 1. Figure S2 for GenBank accession numbers).

#### Development of the ETAS-PCR/Vgsc1014 assay

The ETAS-PCR/*Vgsc*-1014 multiplex-assay is a single PCR reaction, followed by an endonuclease digestion. The strategy for amplification and allelic-discrimination of the three alternative alleles is depicted in Figure 1A.

**Figure 1.**
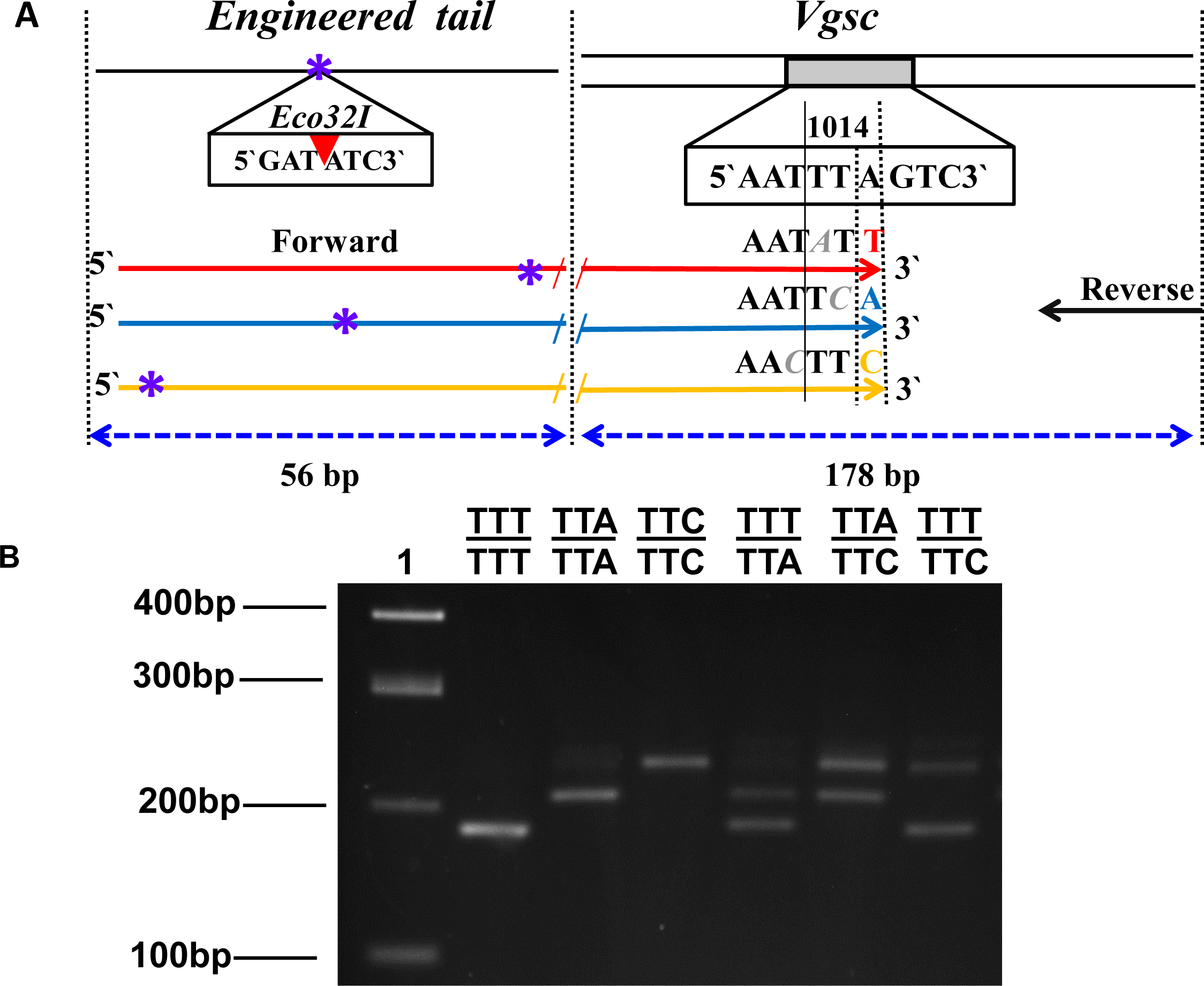
Allele-Specific PCR (AS-PCR) for genotyping tri-allelic *Vgsc*-L1014F variation in *C. quinquefasciatus.* **A.** Schematic representation of the ETAS-PCR/*Vgsc*-1014 assay design with primer locations and predicted size of PCR products. Arrows indicate the location of PCR primers; black – universal reverse primer and red, blue and yellow arrows indicate each allele-specific primer (Codons TTT, TTA and TTC, respectively). Bases highlighted in Grey at the 3’-ends of each specific primer are deliberate mismatches. Blue-asterisks represent the restriction-site of the *Eco*32I enzyme. Full-length primer sequences are reported in Table 1. **B.** An example of the ETAS-PCR/*Vgsc*-1014 gel electrophoresis. Lane 1: 100 bp DNA ladder, lanes 2–4: homozygote for each codon; TTT, TTA and TTC, respectively; lanes 5–6: heterozygous individuals TTT/TTA, TTA/TTC and TTT/TTC, respectively.

To amplify a genomic region of 178 bp encompassing the *Vgsc*-1014 locus by multiplex-PCR reaction, four primers were designed based on a conserved sequence region (Additional file Figure S1) using Primer 3 software (31); with a universal reverse primer and three specific forward primers (Figure 1A). For each allele-specific primer, the 3’-end terminus was designed to match exclusively one of the three SNP alleles (TTA, TTT, TTC) at the last base of the *Vgsc*-1014 codon. To enhance specificity, a deliberate mismatch was also introduced within the last three bases (See Figure 1A).

**Table 1.**
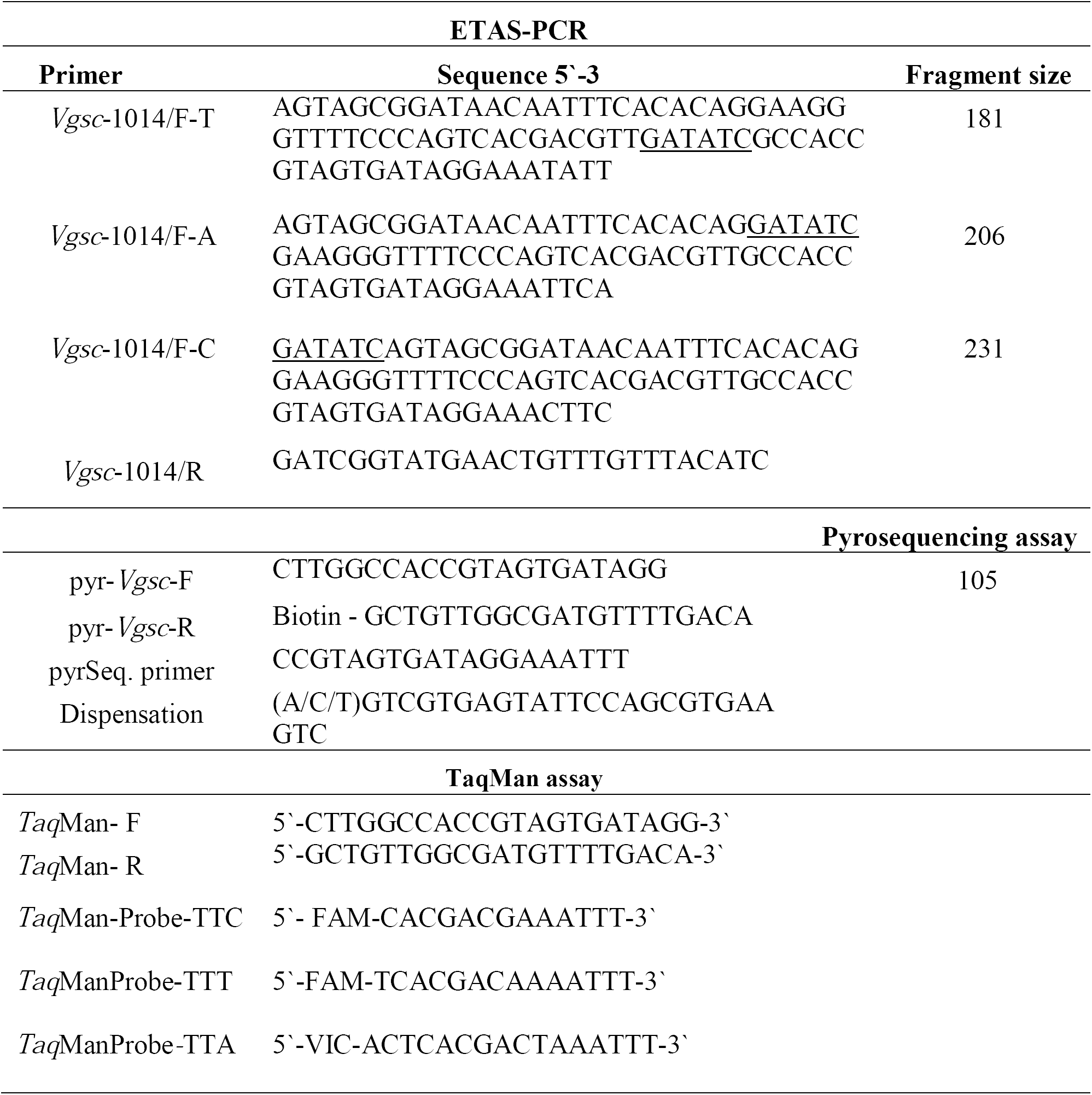
Primers used in the ETAS-PCR/*Vgsc*-1014, pyrosequencing and *Taq*Man assays for genotyping the *Vgsc*-L1014F in *C. quinquefasciatus* mosquitoes. The *Eco*32I site is underlined.

To allow allelic discrimination by agarose gel electrophoresis, three 56bp engineered-tails were synthesized at the 5’ end of the AS forward primer, differing from each other by the engineered position of an *Eco*32I restriction site (5’-GATATC-3’), which was introduced at distinct points of each tail (Figure 1A). Thus, after digestion of the 234bp ETAS-PCR amplified products, each alternative allele has a distinct fragment length (Figure 1B): TTT (181 bp), TTA (206 bp) and TTC (231 bp).

#### ETAS-PCR/Vgsc-1014 amplification, endonuclease digestion and genotyping

The ETAS-PCR/*Vgsc*-1014 multiplex reaction was performed in a total volume of 25 µl containing 10 ng of gDNA, 200 µM each dNTP, 1x PCR buffer, 0.2 units HotStartTaq DNA polymerase (Qiagen), 0.6 µM universal reverse primer *Vgsc*-1014/R, 0.35 µM of the specific forward primers (*Vgsc*-1014/F-T and *Vgsc*-1014/F-C) and 0.3 µM of the *Vgsc*-1014/F-A primer (Table 1). After initial denaturation at 95 °C for 15 min, amplification was performed for 40 cycles of 94°C for 30 s, 59°C for 30 s, and 72°C for 30 s, followed by a final extension step of 72°C for 10 min.

After amplification, the ETAS-PCR/*Vgsc*-1014 multiplex products were digested with FastDigest *Eco*32I restriction enzyme (Thermo Scientific) in a final reaction volume of 30 µl including 10 µl of the PCR product, 2 µl of 10x FastDigest Green buffer, 1.5 µl of FastDigest enzyme and 17 µl distilled water. The reaction was incubated at 37°C for 10 min. Finally, 10 µl of digestion reaction was loaded on a 3% agarose gel.

#### Vgsc-1014 Screening by Pyrosequencing assay

For pyrosequencing genotyping, PCR reactions were performed in a total of 25 µl containing 10 ng of gDNA, 200 µM of each dNTP, 1x PCR buffer, 2.0 mM MgCl_2,_ 0.6 units of HotStartTaq DNA polymerase (Qiagen) and 0.4 µM of primers pyr-*Vgsc*-F and pyr-*Vgsc*-R (Table 1)(21). After initial denaturation at 95°C for 15 min, PCR amplification was performed for 40 cycles of 94°C for 30 s, 58°C for 30 s, and 72°C for 30 s, followed by a final extension step at 72°C for 10 min.

The pyrosequencing genotyping was performed using single-stranded PCR products obtained using the PyroMark Q24 Vacuum Prep Workstation then used in pyrosequencing reactions performed using the PyroMark Gold Q96 reagent kit (Qiagen). The sequencing primer and dispensation order are described in Table 1.

#### Vgsc-1014 genotyping by TaqMan allelic discrimation

Due to the tri-allelic variation in the locus *Vgsc-*1014, two distinct TaqMan assays are applied in parallel for each sample; one to genotype TTA/TTC alleles and the other to detect TTA/TTT variants (27). Primers and TaqMan probes are shown in Table 1.

### Association between target-site alleles and resistant phenotypes

To verify the presence of *Vgs*c-1014 alleles in insecticide exposed mosquitoes, 3-5 day-old adult F1 female mosquitoes from Tororo, Uganda, were exposed to three insecticides using WHO papers impregnated with a diagnostic concentration: DDT (4%), lambda-cyhalothrin (0.05%) or deltamethrin (0.05%). Mosquitoes were exposed to DDT and lambda-cyhalothrin for 1 h and to deltamethrin for 4 h, in four replicates of 25 non-blood fed mosquitoes at an average temperature of 26°C and humidity of 63%. Control bioassays were performed with 25 mosquitoes exposed to non-insecticide treated papers. Insecticide exposed mosquitoes were then transferred to clean holding tubes and provided with 10% glucose for a 24-hour period after which mortality was recorded and alive and dead mosquitoes were collected and individually stored on silica gel.

## Results and Discussion

To determine if the newly-designed ETAS-PCR/*Vgsc*-1014 performed consistently across *C. quinquefasciatus* wild populations, genotyping was conducted on mosquitoes from different continents (Africa and South America). Samples from multiple locations in Africa (*N*=74); Uganda (Tororo and Jinja) and Tanzania (Mwanza) were genotyped by pyrosequencing and 73 (98.6%) agreed with the ETAS-PCR/*Vgsc*-1014 scores, with the single ETAS-PCR discordant result resulting from low PCR efficiency for that sample, which does not allowed detention of PCR fragments after digestion. For the TaqMan assay, mosquitoes from Campina Grande, Brazil (*N*=24) and pyrethroid-exposed individuals from Tororo, Uganda (*N* =84) were genotyped, with 100% agreement to ETAS-PCR/*Vgsc*-1014.

The ETAS-PCR/*Vgsc*-1014 genotyping compares extremely well to the standard assays and represents a more convenient alternative for low technology laboratories because it uses relatively inexpensive laboratory methods (standard PCR and agarose-gel electrophoresis) while can also genotype variation at this tri-allelic SNP in a single reaction. For instance, the genotyping cost for ETAS-PCR is around 4.6 and 3.12 times lower compared to TaqMan and pyrosequencing, respectively, based on the minimum chemistry required (e.g. DNA Polymerase or master mix, either standard or biotinylated primers or fluorescence probes).

The ETAS-PCR/*Vgsc*-1014 also addresses limiations reported in other Allelic-Specific PCR (AS-PCR) assays designed for typing triple variants within a codon. For instance, the AS-PCR applied for genotyping *Vgsc*-L1014F and *Vgsc*-L1014S in *Culex pipiens pallens*, can differentiate wild-type from resistance-alleles but lacks the resolution to discriminate between the two resistance alleles (25). Additionally, the AS-PCR of Chamnanya (32) designed to type *Vgsc*-1014 in *C. quinquefasciatus* differentiates only two (TTA and TTT) of the three alleles but can not detect the TTC resistant allele, which we found to be comon in Tanzania and Uganda (Table 2) and has been reproted elsewhere (29). Indeed, in mosquitoes from Tororo, Uganda we did detect a high frequency of heterozygous wild-type/resistance-alleles among the phenotypically resistant mosquitoes exposed to either deltamethrin, lambda-cyhalothrin or DDT, for which was recorded a frequency of genotypes harboring at least one resistance-allele ranging from 70.37% to 85.78% (Additional file 2. TableS1).

**Table 2.** Specificity of ETAS-PCR *Vgsc*-1014 in comparison to pyrosquencing and TaqMan genotyping

| Country<br>/Location | Brazil |  | Uganda |  |  |  |  |  | Tanzania |  |
| --- | --- | --- | --- | --- | --- | --- | --- | --- | --- | --- |
|  | Campina Grande |  | Tororo<sup>a</sup> |  | Tororo |  | Jinja |  | Mwanza |  |
| Genotype | ETAS-PCR | TaqMan | ETAS-PCR | TaqMan | ETAS-PCR | Pyr | ETAS-PCR | Pyr | ETAS-PCR | Pyr |
| TTA/TTA | 24 | 24 | 23 | 23 | 2 | 1 | 4 | 4 | 3 | 3 |
| TTC/TTC |  |  | 15 | 15 | 10 | 10 | 4 | 4 | 2 | 2 |
| TTT/TTT |  |  | 1 | 1 |  |  | 1 | 1 | 1 | 1 |
| TTA/TTC |  |  | 23 | 23 | 1 | 1 | 9 | 9 | 8 | 8 |
| TTA/TTT |  |  | 9 | 9 |  |  | 3 | 3 | 7 | 7 |
| TTC/TTT |  |  | 13 | 13 | 12 | 13 | 3 | 3 | 2 | 2 |
| **N (98)** |  | **24** |  | **84** |  | **28** |  | **24** |  | **23** |

<sup>a</sup> mosquito exposed to insecticides (WHO - Tube Bioassay)
Pyr; Pyrosequencing

## Conclusion

Therefore, since the evolution of insecticide resistance could increase the transmission burden for all major neglected tropical diseases, developing tools which are easy to apply, such as the ETAS-PCR/*Vgsc*-1014, is imperative to support decision-making for resistance management and to preserve the effectiveness of the restricted number of insecticides available for vector control.

## Abbreviations

LF: lymphatic filariasis
*Vgsc*: voltage gated sodium channel
ETAS-PCR: Engineered-Tail Allele-Specific-PCR
WNV: West Nile Virus
SLEV: St. Louis encephalitis Virus
MDA: mass drug administration
AS-PCR: Allelic-Specific PCR

## Declarations

### Ethics approval and consent to participate

Not applicable

### Consent for publication

Not applicable

### Availability of data and materials

All data generated or analysed during this study are included in this published article and its supplementary information files

## Contributions

WFSM. Conceived, designed the experiments and wrote the paper. WFSM, BNSP, ATVA, AM, and PGSM. performed the experiments. WFSM. and BNSP analysed the data. BNSP, ATVA, AM, P.S.M, DW and MJD. Contributed reagents/materials/sample collections. DW, CSW. and MJD. reviewed the final manuscript. All authors read and approved the final manuscript.

## Acknowledgments

We would like to thank Henry Mawejje for supporting the field-collections in Uganda. The authors also thank Dr. Triantafillos Liloglou and Keith Steen for supporting the pyrosequencing genotyping.

## Additional files

**Additional file 1: Figure S1**. Multiple sequences alignment of the partial fragment of the *Vgsc* gene across *Culex pipiens pipiens, Culex p. quinquefasciatus* and *Culex p. pallens*. **Figure S2**. Neighbor-joining tree of the partial fragment of the *Vgsc* gene.

Additional file 2: **Table S1.** Allelic and genotypic frequency of *Vgsc*-1014F in relation to mosquito survival phenotype by deltamethrin, lambda-cyhalothrin or DDT in *C. quinquefasciatus* from Tororo, Uganda.

