## additional File 1 for "Development and Application of a Tri-allelic PCR Assay for Screening *Vgsc*-L1014F *Kdr* Mutations Associated with Pyrethroid and Organochlorine Resistance in the Mosquito *Culex quinquefasciatus*"

**Forward**

```
C.quinq/KR061930.1/1-305      1 GTGGGCGACGTGTCCTGCATTCCGTTCTTCTTGGCCACCGTAGTGATAGGA 51
C.quinq/KR061940.1/1-307      1 GTGGGCGACGTGTCCTGCATTCCGTTCTTCTTGGCCACCGTAGTGATAGGA 51
C.quinq/KR061970.1/1-306      1 GTGGGCGACGTGTCCTGCATTCCGTTCTTCTTGGCCACCGTAGTGATAGGA 51
C.quinq/KM377241.1/1-307      1 GTGGGCGACGTGTCCTGCATTCCGTTCTTCTTGGCCACCGTAGTGATAGGA 51
C.quinq/KM377242.1/1-307      1 GTGGGCGACGTGTCCTGCATTCCGTTCTTCTTGGCCACCGTAGTGATAGGA 51
C.quinq/EF658661.1/1-307      1 GTGGGCGACGTGTCCTGCATTCCGTTCTTCTTGGCCACCGTAGTGATAGGA 51
C.quinq/EF658660.1/1-305      1 GTGGGCGACGTGTCCTGCATTCCGTTCTTCTTGGCCACCGTAGTGATAGGA 51
C.quinq/EF658659.1/1-305      1 GTGGGCGACGTGTCCTGCATTCCGTTCTTCTTGGCCACCGTAGTGATAGGA 51
C.quinq/DQ497226.1/1-306      1 GTGGGCGACGTGTCCTGCATTCCGTTCTTCTTGGCCACCGTAGTGATAGGA 51
C.quinq/DQ497227.1/1-306      1 GTGGGCGACGTGTCCTGCATTCCGTTCTTCTTGGCCACCGTAGTGATAGGA 51
C.pallens/GU198930.1/1-307    1 GTGGGCGACGTGTCCTGCATTCCGTTCTTCTTGGCCACCGTAGTGATAGGA 51
C.pallens/HQ540623.1/1-307    1 GTGGGCGACGTGTCCTGCATTCCGTTCTTCTTGGCCACCGTAGTGATAGGA 51
C.pipiens/DQ497213.1/1-307    1 GTGGGCGACGTGTCCTGCATTCCGTTCTTCTTGGCCACCGTAGTGATAGGA 51
C.pipiens/DQ497230.1/1-306    1 GTGGGCGACGTGTCCTGCATTCCGTTCTTCTTGGCCACCGTAGTGATAGGA 51
C.pipiens/DQ497228.1/1-306    1 GTGGGCGACGTGTCCTGCATTCCGTTCTTCTTGGCCACCGTAGTGATAGGA 51
C.pipiens/DQ497214.1/1-306    1 GTGGGCGACGTGTCCTGCATTCCGTTCTTCTTGGCCACCGTAGTGATGGGA 51

C.quinq/KR061930.1/1-305     52 AATTTAGTCGTGAGTATTCCAGCGTGAAGTCTTAGCGATTGATCTAGTGTG 102
C.quinq/KR061940.1/1-307     52 AATTTTGTCGTGAGTATTCCAGCGTGAAGTCTTAGCGATTGATCTAGTGTG 102
C.quinq/KR061970.1/1-306     52 AATTTCGTCGTGAGTATTCCAGCGTGAAGTCTTCGTGATTGATCTAGTGTG 102
C.quinq/KM377241.1/1-307     52 AATTTTGTCGTGAGTATTCCAGCGTGAAGTCTTAGCGATTGATCTAGTGTG 102
C.quinq/KM377242.1/1-307     52 AATTTAGTCGTGAGTATTCCAGCGTGAAGTCTTAGCGATTGATCTAGTGTG 102
C.quinq/EF658661.1/1-307     52 AATTTTGTCGTGAGTATTCCAGCGTGAAGTCTTAGCGATTGATCTAGTGTG 102
C.quinq/EF658660.1/1-305     52 AATTTAGTCGTGAGTATTCCAGCGTGAAGTCTTAGCGATTGATCTAGTGTG 102
C.quinq/EF658659.1/1-305     52 AATTTAGTCGTGAGTATTCCAGCGTGAAGTCTTAGCGATTGATCTAGTGTG 102
C.quinq/DQ497226.1/1-306     52 AATTTTGTCGTGAGTATTCCAGCGTGAAGTCTTAGCGATTGATCTAGTGTG 102
C.quinq/DQ497227.1/1-306     52 AATTCAGTCGTGAGTATTCCAGCGTGAAGTCTTAGCGATTGATCTAGTGTG 102
C.pallens/GU198930.1/1-307   52 AATTTTGTCGTGAGTATTCCAGCGTGAAGTCTTAGCGATTGATCTAGTGTG 102
C.pallens/HQ540623.1/1-307   52 AATTTTGTCGTGAGTATTCCAGCGTGAAGTCTTAGCGATTGATCTAGTGTG 102
C.pipiens/DQ497213.1/1-307   52 AATTTTGTCGTGAGTATTCCAGCGTGAAGTCTTAGCGATTGATCTAGTGTG 102
C.pipiens/DQ497230.1/1-306   52 AATTTAGTCGTGAGTATTCCAGCGTGAAGTCTTAGCGATTGATCTAGTGTG 102
C.pipiens/DQ497228.1/1-306   52 AATTTAGTCGTGAGTATTCCAGCGTGAAGTCTTCGTGATTGATCTAGTGTG 102
C.pipiens/DQ497214.1/1-306   52 AATTTTGTCGTGAGTATTCCAGCGTGAAGTCTTCGTGATTGATCTAGTGTG 102

C.quinq/KR061930.1/1-305    103 CGCGCTAGAGCTGTCAAAACATCGCCAACAGCATGCAAGAAAAGGTGGGAA 153
C.quinq/KR061940.1/1-307    103 CGCGCTAGAGCTGTCAAAACATCGCCAACAGCATGCAAGAAAAGGTGGGAA 153
C.quinq/KR061970.1/1-306    103 CGCATTAGAGCTGTCAAAACATCGCCAACAGCATGCAAGAAAAGGTGGGAA 153
C.quinq/KM377241.1/1-307    103 CGCGCTAGAGCTGTCAAAACATCGCCAACAGCATGCAAGAAAAGGTGGGAA 153
C.quinq/KM377242.1/1-307    103 CGCGCTAGAGCTGTCAAAACATCGCCAACAGCATGCAAGAAAAGGTGGGAA 153
C.quinq/EF658661.1/1-307    103 CGCGCTAGAGCTGTCAAAACATCGCCAACAGCATGCAAGAAAAGGTGGGAA 153
```

| | | | |
|---|---|---|---|
| *C.quinq/KR061930.1/1-305* | 103 | CGCGCTAGAGCTGTCAAAACATCGCCAACAGCATGCAAGAAAAGGTGGGAA | 153 |
| *C.quinq/KR061940.1/1-307* | 103 | CGCGCTAGAGCTGTCAAAACATCGCCAACAGCATGCAAGAAAAGGTGGGAA | 153 |
| *C.quinq/KR061970.1/1-306* | 103 | CGCATTAGAGCTGTCAAAACATCGCCAACAGCATGCAAGAAAAGGTGGGAA | 153 |
| *C.quinq/KM377241.1/1-307* | 103 | CGCGCTAGAGCTGTCAAAACATCGCCAACAGCATGCAAGAAAAGGTGGGAA | 153 |
| *C.quinq/KM377242.1/1-307* | 103 | CGCGCTAGAGCTGTCAAAACATCGCCAACAGCATGCAAGAAAAGGTGGGAA | 153 |
| *C.quinq/EF658661.1/1-307* | 103 | CGCGCTAGAGCTGTCAAAACATCGCCAACAGCATGCAAGAAAAGGTGGGAA | 153 |
| *C.quinq/EF658660.1/1-305* | 103 | CGCGCTAGAGCTGTCAAAACATCGCCAACAGCATGCAAGAAAAGGTGGGAA | 153 |
| *C.quinq/EF658659.1/1-305* | 103 | CGCGCTAGAGCTGTCAAAACATCGCCAACAGCATGCAAGAAAAGGTGGGAA | 153 |
| *C.quinq/DQ497226.1/1-306* | 103 | CGCGCTAGAGCTGTCAAAACATCGCCAACAGCATGCAAGAAAAGGTGGGAA | 153 |
| *C.quinq/DQ497227.1/1-306* | 103 | CGCGCTAGAGCTGTCAAAACATCGCCAACAGCATGCAAGAAAAGGTGGGAA | 153 |
| *C.pallens/GU198930.1/1-307* | 103 | CGCGCTAGAGCTGTCAAAACATCGCCAACAGCATGCAAGAAAAGGTGGGAA | 153 |
| *C.pallens/HQ540623.1/1-307* | 103 | CGCGCTAGAGCTGTCAAAACATCGCCAACAGCATGCAAGAAAAGGTGGGAA | 153 |
| *C.pipiens/DQ497213.1/1-307* | 103 | CGCGCTGGAGCTGTCAAAACATCGCCAACAGCATGCAAGAAAAGGTGGGAA | 153 |
| *C.pipiens/DQ497230.1/1-306* | 103 | CGCGCTAGAGCTGTCAAAACATCGCCAACAGCATGCAAGAAAAGGTGGGAA | 153 |
| *C.pipiens/DQ497228.1/1-306* | 103 | CGCATTAGAGCTGTCAAAACATCGCCAACAGCATGCAAGAAAAGGTGGGAA | 153 |
| *C.pipiens/DQ497214.1/1-306* | 103 | CGCATTAGAGCTGTCAAAACATCGCCAACAGCATGCAAGAAAAGGTGGGAA | 153 |

**Reverse**

| | | | |
|---|---|---|---|
| *C.quinq/KR061930.1/1-305* | 154 | CGAAAAACTTTAAGGTCACATTTGTACCTTTGATGTAAACAAACAGTTCAT | 204 |
| *C.quinq/KR061940.1/1-307* | 154 | CGAAAAACTTTAAGGTCACATTTGTACCTTTGATGTAAACAAACAGTTCAT | 204 |
| *C.quinq/KR061970.1/1-306* | 154 | CGAAAAACTTTAAGGTCACATTTGTACCTTTGATGTAAACAAACAGTTCAT | 204 |
| *C.quinq/KM377241.1/1-307* | 154 | CGAAAAACTTTAAGGTCACATTTGTACCTTTGATGTAAACAAACAGTTCAT | 204 |
| *C.quinq/KM377242.1/1-307* | 154 | CGAAAAACTTTAAGGTCACATTTGTACCTTTGATGTAAACAAACAGTTCAT | 204 |
| *C.quinq/EF658661.1/1-307* | 154 | CGAAAAACTTTAAGGTCACATTTGTACCTTTGATGTAAACAAACAGTTCAT | 204 |
| *C.quinq/EF658660.1/1-305* | 154 | CGAAAAACTTTAAGGTCACATTTGTACCTTTGATGTAAACAAACAGTTCAT | 204 |
| *C.quinq/EF658659.1/1-305* | 154 | CGAAAAACTTTAAGGTCACATTTGTACCTTTGATGTAAACAAACAGTTCAT | 204 |
| *C.quinq/DQ497226.1/1-306* | 154 | CGAAAAACTTTAAGGTCACATTTGTACCTTTGATGTAAACAAACAGTTCAT | 204 |
| *C.quinq/DQ497227.1/1-306* | 154 | CGAAAAACTTTAAGGTCACATTTGTACCTTTGATGTAAACAAACAGTTCAT | 204 |
| *C.pallens/GU198930.1/1-307* | 154 | CGAAAAACTTTAAGGTCACATTTGTACCTTTGATGTAAACAAACAGTTCAT | 204 |
| *C.pallens/HQ540623.1/1-307* | 154 | CGAAAAACTTTAAGGTCACATTTGTACCTTTGATGTAAACAAACAGTTCAT | 204 |
| *C.pipiens/DQ497213.1/1-307* | 154 | CGAAAAACTTTAAGGTCACATTTGTACCTTTGATGTAAACAAACAGTTCAT | 204 |
| *C.pipiens/DQ497230.1/1-306* | 154 | CGAAAAACTTTAAGGTCACATTTGTACCTTTGATGTAAACAAACAGTTCAT | 204 |
| *C.pipiens/DQ497228.1/1-306* | 154 | CGAAAAACTTTAAGGTCACATTTGTACCTTTGATGTAAACAAACAGTTCAT | 204 |
| *C.pipiens/DQ497214.1/1-306* | 154 | CGAAAAACTTTAAGGTCACATTTGTACCTTTGATGTAAACAAACAGTTCAT | 204 |

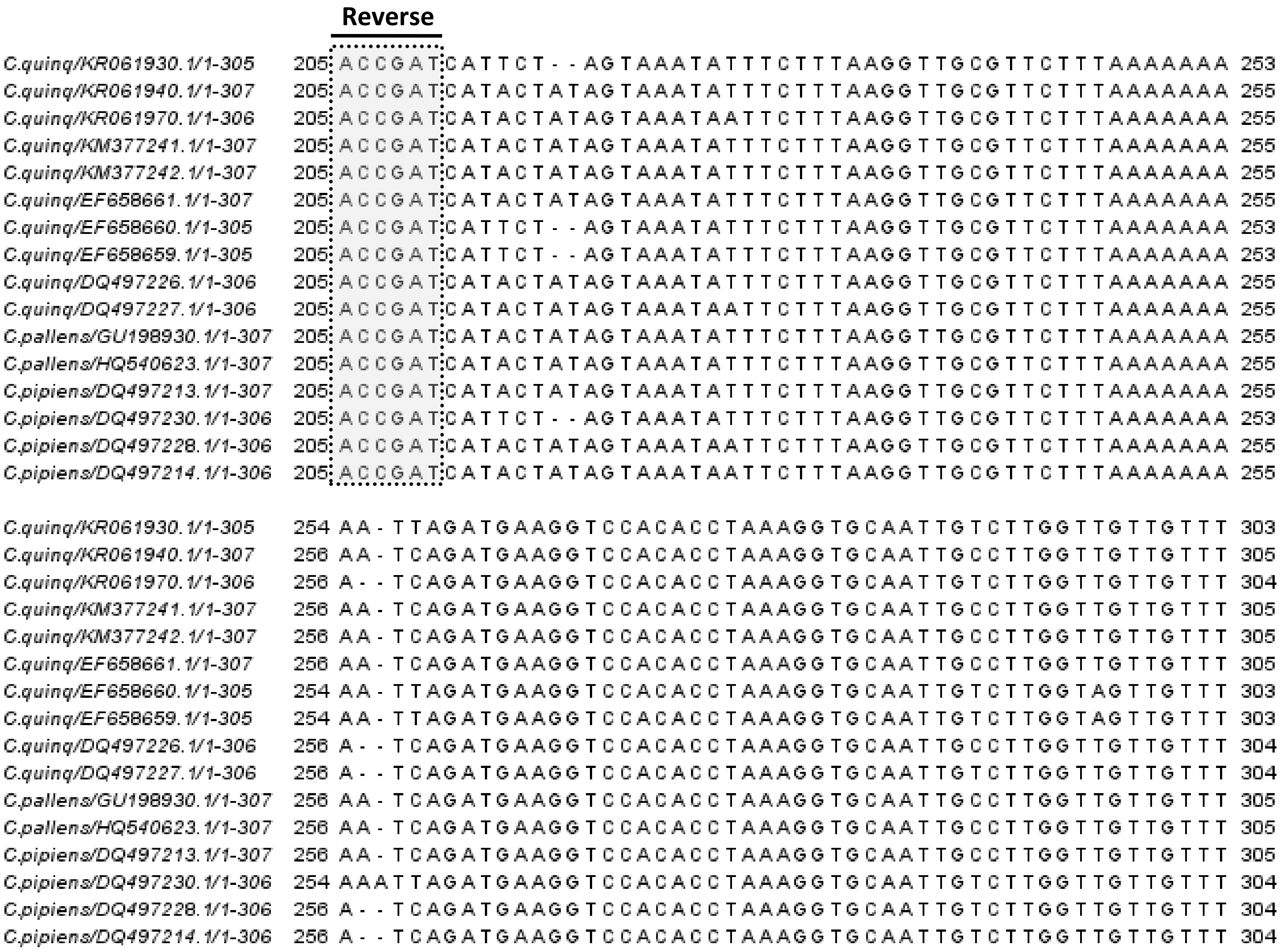


**Figure S1.** Multiple sequences alignment of the partial fragment of the *Vgsc* gene across *Culex pipiens pipiens*, *Culex p. quinquefasciatus* and *Culex p. pallens*. Alignment was performed using MEGA software (1). Grey shaded boxes correspond to forward and reverse primer sequences. The red line in the forward primer highlights the position of the *Vgsc*-1014 mutation.

1. Tamura K, Peterson D, Peterson N, Stecher G, Nei M, Kumar S. MEGA5: molecular evolutionary genetics analysis using maximum likelihood, evolutionary distance, and maximum parsimony methods. Molecular Biology and Evolution. 2011;28(10):2731-9.

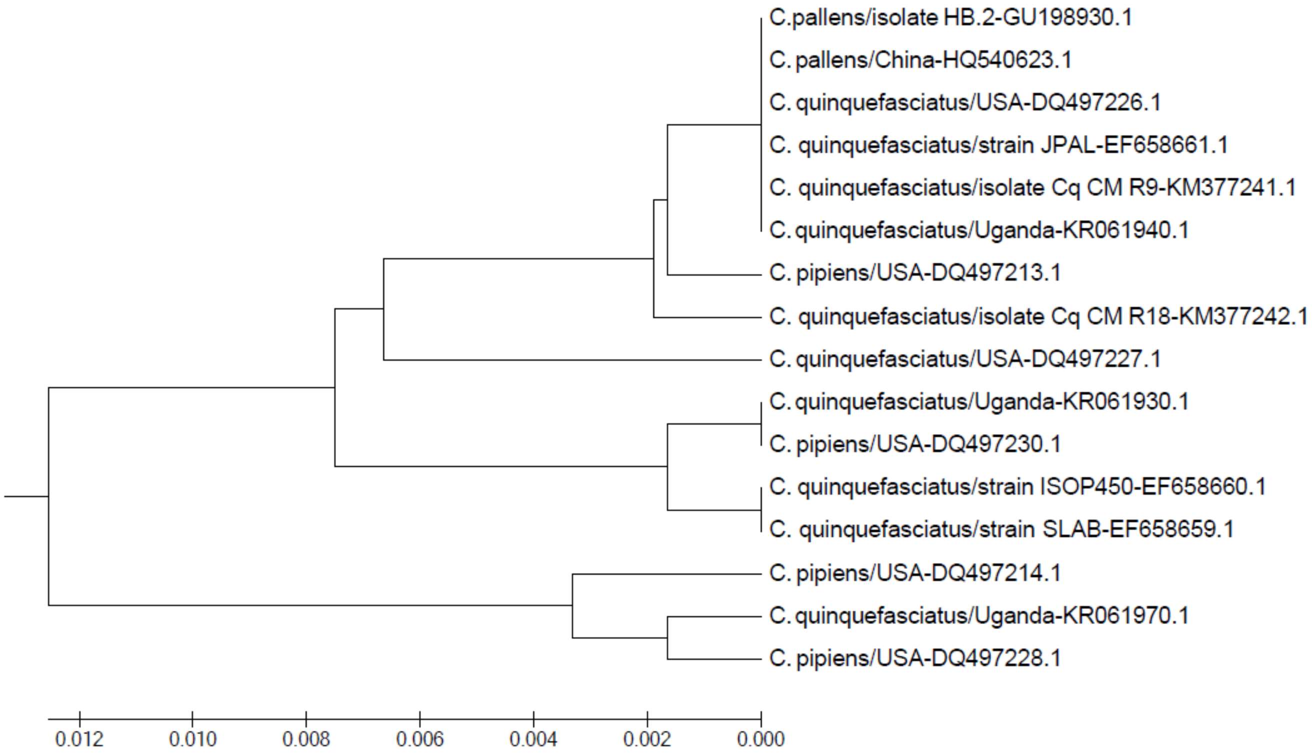


**Figure S2.** Neighbor-joining tree of the partial fragment of the *Vgsc* gene. Branch names are composed of; *Culex* species/ geographic origin or strain name, and GenBank accession number.
