## Additional file 2 for "Development and Application of a Tri-allelic PCR Assay for Screening *Vgsc*-L1014F *Kdr* Mutations Associated with Pyrethroid and Organochlorine Resistance in the Mosquito *Culex quinquefasciatus*"

**Table S1.** Allelic and genotypic frequency of *Vgsc*-1014F in relation to mosquito survival phenotype by deltamethrin, lambda-cyhalothrin or DDT in *C. quinquefasciatus* from Tororo, Uganda.

| Insecticide | N | Survivorship | Phenotype | Genotype | | | | | | *Vgsc*-1014F allele freq[a] | *Vgsc*-1014F genotype freq[b] |
|---|---|---|---|---|---|---|---|---|---|---|---|
| | | | | AA** | CC | TT | AC | AT | CT | | |
| Deltamethrin | 24 | 66.66% *(44.66-83.66) | Alive | 3 | 7 | | 5 | | 1 | 65.6% | 81.25 |
| | | | Dead | 3 | 2 | | 1 | 1 | 1 | | |
| lambda-cyhalothrin | 30 | 93.33% *(77.9-99.13) | Alive | 4 | 1 | | 8 | 7 | 8 | 58.8% | 85.71 |
| | | | Dead | 2 | | | | | | | |
| DDT | 30 | 90% *(73.5-97.9) | Alive | 8 | 5 | 1 | 9 | 1 | 3 | 51.8% | 70.37 |
| | | | Dead | 3 | | | | | | | |

* 95% confidence intervals

** Wild-type genotype

[a]; frequency within alive

[b]; frequency within alive excluding wild-type homozygous
